# Antibiotic-resistant bacteria in agricultural soil appraised against non-agricultural soil: case in one of the most agriculturally-active settings in West Cameroon

**DOI:** 10.1101/2024.08.27.609998

**Authors:** Loïc Kevin Kamgaing Nkamguia, O’Neal Dorsel Youté, Blandine Pulcherie Tamatcho Kweyang, Esther Guladys Kougang, Pascal Blaise Well a Well a Koul, Adolarice Nana Feukeu, Christelle Domngang Noche, Pierre René Fotsing Kwetche

## Abstract

**Background:** Agriculture is also concerned by the problem of bacterial resistance because agricultural soils are reservoirs of antibiotic-resistant bacteria (ARB) and antibiotic-resistant genes.

**Objective:** An investigation about ARB was carried out on agricultural farms soils in Mangoum, a neighborhood of the Foumbot municipality (Noun division, West Cameroon).

**Method:** It was conducted as a cross-sectional descriptive study with a total of 46 soil specimens collected from plant farms and a control plot. Isolation, enumeration and antibiotic susceptibility tests were performed according to standard protocols.

**Results:** The bacteria recovered basically included *Aeromonas* spp., *Chryseobacterium* spp., *Pseudomonas* spp., *Staphylococcus* spp., and Gram-positive rods. Their loads in the farmland soils were significantly lower than in the control plot. Overall, susceptibility tests performed with 169 bacterial colony morphotypes revealed high resistant rates. Also, most of isolates expressed multidrug-resistance to the antibiotics used, while highest resistance rates were recorded with isolates form agricultural plots. Levofloxacin, Imipenem, Gentamicin and Ciprofloxacin were globally the most effective.

**Conclusion:** Agreeing with previous surveys conducted on animal farms with similar environmental conditions, these findings could provide support to sustainable orientation of policies regarding control of antimicrobial resistance in Cameroon from the One Health perspective.

## Introduction

One of the most significant progresses made in health throughout the 20^th^ century was the development of antimicrobial agents used to control microorganisms which are often etiologies of health disorders in humans, animals and plants. These advances in health-related microbiology resulted in major achievements in connection with alleviated morbidity and mortality due to infectious diseases across global human populations. The recorded achievements and further anticipations were, however, undermined by microbial resistance expressed against previously effective drugs [1-3]. The emergence and spread of antibiotic-resistance bacteria (ARB) is currently known to result from selective pressure exerted by diverse engines like antibacterial agents or other antimicrobials that promote activity of mobile genetic determinants which carry resistance traits, in order to support population fitness in adverse environmental conditions [4,5].

Agricultural soils are known reservoirs for ARB and antibiotic resistant genes (ARG) [6,7]. Some mobiles recognized as factors for selection and dissemination of resistance traits in plant farms include the use of pesticides and manure in crop production [3,5], though their contribution is yet to be accurately assessed because of the versatility in compositions and application. In fact, antibiotics used as food supplements in farm animals is consistently pointed out as a major cause of selection for resistant phenotypes that diffuse throughout diverse close and remote bacterial populations as well as animal feces, vehicle of residual or non-metabolized antibiotics in exposed environments. Consequently, since animal manure (made from animal feces) serves as soils fertilizers in crop production, the likelihood for selection and spread of ARB and ARG in vulnerable environmental settings is high [8]. Like other selective agents, biocides used to improve agricultural yields can also cause pressure that co-select antibiotic-resistant bacterial populations [9,10]. Selection in all environments is facilitated by the high flexibility of soil prokaryotic genomes that explain their rapid adaptation to new environments, and to the newly introduced chemical compounds [5].

These pieces of information support all investigations in the framework of AMR in agriculture, acknowledging that altered microbial populations will not only affect the quality of soils and the quality of crop that can effectively grow, but also exposed human and animals as well. In a lager project aiming at assessing AMR and contributing factors, the present investigation was conducted in farmland soils of one amongst the most important crop production basin in West Cameroon. Together with related findings in animal and plant farms, recovered information will guide decision makers in developing suitable policies that will advocate and support sustainably the antimicrobial resistance stewardship in plant farms, aligning with those in connection with human and animal health according to the One Health principles.

## Material and methods

### Study design and ethical/administrative considerations

Data collection in the present cross-sectional study was conducted in agricultural farms located in Mangoum, a neighborhood of the Foumbot municipality (Noun division in West Cameroon). Thereafter, specimen analyses were performed in the Laboratory of Microbiology at “Université des Montagnes” Teaching Hospital (UdMTH). This work was conducted between September and October, 2020.

Before field works, administrative approvals were obtained from legal authorities. Authorizations were provided by the Foumbot sub-divisional officer and the division Head for the Ministry of Agriculture and Rural Development. Also, prior to field sample collection, all farm owners signed a voluntary written informed consent to authorize the research to be carried out on their lands. For the present study, ethical approval was not required because humans were not the subjects of interest. The work was carried out only on environmental samples, particularly soil specimens; farmers were not at risk.

Laboratory screening was performed under authorization reference N° 2020/176/AED/UDM/CUM delivered by the UdMTH General Administration. Specimen collection was performed in the farms in which the owner was affiliated and recorded in the local divisional headquarter for the Ministry of Agriculture and Rural Development.

### Data collection

An adapted data collection sheet was used to gather relevant pieces of information in connection with the investigation goals. These pieces of information included type of manure used, source of manure, frequency of manure use, most recent date of manure application, pesticide use, frequency of pesticide use, biosecurity-biosafety practices, and training in agricultural activities.

### Samples collection and transport

In line with bio-safety and biosecurity rules, portions of approximately 50 g of soil were randomly collected with sterile spatula at different locations in each farm, and preserved separately in labelled sterile containers. The same procedure was used to collect soil specimens from a nearby virgin piece of land (referred as a “control plot”) with wild vegetation, no visible chemical or manure application or no other visible anthropogenic activity. Relatively close to each other, all the farms and the control plot were located in the same area. Samples were stored in refrigerated containers and conveyed to the laboratory for screening that was performed within the 24 hours post-collection.

### Bacterial culture, enumeration and identification

After sampling, culture, enumeration and identification of bacteria were performed in line with previous protocols [11].

#### Culture

In the laboratory, each portion of soil was thoroughly mixed to homogeneity. Then, 5 g of the resulting preparation was added to, and thoroughly mixed with 45 mL of sterile physiological saline. Thereafter, a series of successive decimal dilutions were made in sterile physiological saline. From each suspension (diluted and undiluted), 50 μL of the inoculum was spread over the entire surface of McConKey and Mannitol Salt isolation agars with a sterile Pasteur pipette rake. The inoculated agar plates were then incubated overnight (for 24 hours) at 37°C.

#### Identification I: Macroscopy and enumeration

After incubation, bacterial growth on each culture medium was assessed and those on which bacterial growth was recorded were characterized. The emerging characteristics included colonies description (according to their shape, size, consistency, color, surface and opacity). According to morphotypes, differential enumeration of the colonies was carried out. Only plates on which colony count varied between 30 and 300 were used in this exercise. For each soil sample, the bacterial load (*N*) expressed in terms of Colony Forming Units per gram of soil (CFU/g of soil) was calculated according to the formula *N* = 900 × *p* × 10^*d*^ / 5 (where 900-is the ratio of the volume of the initial suspension to the volume of the inoculum, *p*-the number of Colony Forming Units (CFU) per Petri dish, 10^*d*^ -the dilution factor, *d*-the dilution number, 1/5-the conversion factor from the number of CFU in 5 g of soil sample to the number of CFU in 1 g of soil).

#### Identification II: microscopy and bio-enzymatic identification

Subsequent to enumeration, bio-enzymatic identification steps were conducted on distinct colony types. Gram stain was then followed by exploration of bacterial metabolism according to target bacterial group. This exploration was carried according to standard procedures with a series of identification parameters including the oxidase test, tests on Kligler Hajna agar (glucose and lactose fermentation, gas and hydrogen sulfide production), tests on Mannitol-Mobility medium (mannitol fermentation, bacteria mobility, nitrate reductase production), urease, indole, TDA, gelatinase, Voges-Prokauer, decarboxylase (ADH, ODC, LDC), catalase, free coagulase and DNAase tests for Gram-negative rods and Gram-positive cocci, according to basic recommended principles. Concerning Gram-positive rods, identification was limited to microscopy on Gram-stained smear.

### Antibiotic susceptibility tests

Susceptibility tests were carried out by standard disk diffusion according to the “Comité de l’Antibiogramme de la Société Française de Microbiologie, EUCAST” (CASFM) [12]. All tests were conducted with 24 h pure subculture grown on nutritive agar from colonies that were randomly isolated as representative of each colony morphotype in all samples. In total, 14 antibacterial agents commonly used in bacterial infection control in Cameroon were used in subjected bacterial pools. Namely, they were Amoxicillin (20 or 25 μg), Amoxicillin/Clavulanic acid (20/10 μg), Aztreonam (30 μg), Cefepime (30 μg), Cefoxitin (30 μg), Ceftriaxone (30 μg), Ciprofloxacin (5 μg), Gentamicin (10 μg), Imipenem (10 μg), Levofloxacin (5 μg), Norfloxacin (10 μg), Penicillin G (10 U), Tetracycline (30 μg), Trimethoprim/Sulphamethoxazol (1.75/23.25 μg). For the clinical categorization of GPR isolates and Penicillin G (10 U) testing, the 2013 recommendation of CASFM was used [13]. *Staphylococcus aureus* ATCC 29213 and *Escherichia coli* ATCC 25922 were the reference bacterial strain used for quality control throughout the process.

### Data analysis

Data recorded included pieces of information collected from farm owners, types of bacteria recovered, bacterial loads and clinical categories (susceptible-susceptible at high posology-resistant) of studied isolates. These data were recorded and processed with Microsoft Excel 2013 and analyzed with tools provided by IBM SPSS statistic version 20. In this paper, bacterial loads were presented for each sample. About the clinical categories, results were presented in terms of frequencies per bacterial types and antibacterial agents. Linear regression tests were performed to assess association between bacterial loads with the characteristic of “agricultural plot” (that is plot undergoing agricultural transformations and treatments). Significant results were admitted for p-values less than 0.05.

## Results

### Survey results

In this work, six agricultural farms were enrolled. Data analysis from the survey sheet revealed that all farm owners have been trained in agriculture. However, there was a lack of biosafety and biosecurity practices (partial hygiene, which was limited to the hand washing after farming) and an absence of knowledge related with antibiotic resistance. Pesticides and animal manure were used on all farms. From swine and poultry farms, manure was spread before plowing and seeding. The latest application was done between 2 and 6 weeks (depending of the farm) before the present study was initiated. The pesticide application was carried out at 3, 4 or 5 -day intervals (in 3, 1, 2 farms, respectively). No specific reference existed to guide farmers’ practices. Gloves were rarely used if ever, boots also.

### Bacterial diversity and loads

From a total of 46 soil samples collected (Control plot: 5, Farm 1: 5, Farm 2: 6, Farm 3: 6, Farm 4: 8, Farm 5: 9, Farm 6: 7), bacteria recovered included *Aeromonas* spp., *Chryseobacterium* spp., *Pseudomonas* spp., *Staphylococcus* spp., and Gram-positive rods (GPR). *Chryseobacterium* spp., were the least frequently isolated. In addition, bacterial loads in farms specimens were significantly lower than those recorded from the control plot specimens (Tables 1 and 2). Highest bacterial densities were obtained with Gram-positive rods, *Aeromonas* and *Pseudomonas* (Table 2). The linear regression tests indicated that the total bacterial loads and those for each bacterial group were associated with the characteristic “agricultural plot” of farms (for total bacterial loads: p-value <0.001; for *Staphylococcus* spp. and *Pseudomonas* spp. loads: p-value = 0.004; for *Aeromonas* spp. and GPR loads: p-value <0.001).

**Table 1.** Total loads (CFU/g of soil) of bacteria.

| Samples | Control plot | Farm 1 | Farm 2 | Farm 3 | Farm 4 | Farm 5 | Farm 6 |
| --- | --- | --- | --- | --- | --- | --- | --- |
| Sample 1 | 1.0062×10<sup>7</sup> | 2.178×10<sup>4</sup> | 1.656×10<sup>4</sup> | 2.34×10<sup>4</sup> | 1.6452×10<sup>5</sup> | 2.5488×10<sup>6</sup> | 3.7548×10<sup>5</sup> |
| Sample 2 | 3.888×10<sup>7</sup> | 2.934×10<sup>4</sup> | 5.598×10<sup>4</sup> | 3.492×10<sup>4</sup> | 8.802×10<sup>4</sup> | 2.3454×10<sup>6</sup> | 4.14×10<sup>4</sup> |
| Sample 3 | 6.84×10<sup>6</sup> | 1.8162×10<sup>5</sup> | 5.328×10<sup>5</sup> | 4.662×10<sup>4</sup> | 1.5282×10<sup>6</sup> | 2.106×10<sup>4</sup> | 1.3608×10<sup>5</sup> |
| Sample 4 | 4.662×10<sup>6</sup> | 4.374×10<sup>4</sup> | 1.206×10<sup>5</sup> | 3.78×10<sup>6</sup> | 8.568×10<sup>5</sup> | 2.7432×10×10<sup>6</sup> | 1.2942×10<sup>5</sup> |
| Sample 5 | 7.704×10<sup>6</sup> | 2.538×10<sup>4</sup> | 2.556×10<sup>4</sup> | 5.526×10<sup>4</sup> | 1.557×10<sup>5</sup> | 2.6586×10<sup>6</sup> | 2.934×10<sup>4</sup> |
| Sample 6 | - | - | 1.026×10<sup>4</sup> | 5.922×10<sup>4</sup> | 4.41×10<sup>4</sup> | 5.922×10<sup>4</sup> | 4.446×10<sup>4</sup> |
| Sample 7 | - | - | - | - | 2.916×10<sup>4</sup> | 4.392×10<sup>4</sup> | 1.251×10<sup>5</sup> |
| Sample 8 | - | - | - | - | 1.05732×10<sup>6</sup> | 6.498×10<sup>4</sup> | - |
| Sample 9 | - | - | - | - | - | 1.5012×10<sup>6</sup> | - |

CFU: Colony Forming Units, -: absence of visible bacterial growth

**Table 2.** Loads (CFU/g of soil) of various bacterial groups.

| Samples | Control plot | Farm 1 | Farm 2 | Farm 3 | Farm 4 | Farm 5 | Farm 6 | Control plot | Farm 1 | Farm 2 | Farm 3 | Farm 4 | Farm 5 | Farm 6 |
| --- | --- | --- | --- | --- | --- | --- | --- | --- | --- | --- | --- | --- | --- | --- |
|  | *Aeromonas* spp. |  |  |  |  |  |  | *Pseudomonas* spp. |  |  |  |  |  |  |
| Sample 1 | 8.496×10<sup>6</sup> | 7.56×10<sup>3</sup> | - | 3.96×10<sup>3</sup> | - | - | 4.878×10<sup>4</sup> | 1.152×10<sup>6</sup> | - | - | 3.24×10<sup>3</sup> | 1.3176×10<sup>5</sup> | - | 1.656×10<sup>4</sup> |
| Sample 2 | 1.476×10<sup>7</sup> | 1.062×10<sup>4</sup> | 2.16×10<sup>3</sup> | 8.28×10<sup>3</sup> | 1.8×10<sup>4</sup> | - | 1.656×10<sup>4</sup> | 2.25×10<sup>7</sup> | 1.8×10<sup>3</sup> | - | 1.134×10<sup>4</sup> | 5.04×10<sup>3</sup> | 4.878×10<sup>5</sup> | 2.484×10<sup>4</sup> |
| Sample 3 | 5.76×10<sup>5</sup> | 1.008×10<sup>5</sup> | 4.86×10<sup>5</sup> | 9.72×10<sup>3</sup> | 2.232×10<sup>5</sup> | 2.106×10<sup>4</sup> | 1.368×10<sup>4</sup> | 3.78×10<sup>5</sup> | - | - | 1.116×10<sup>4</sup> | - | - | 4.104×10<sup>4</sup> |
| Sample 4 | 1.314×10<sup>6</sup> | 1.494×10<sup>4</sup> | 5.22×10<sup>4</sup> | 8.424×10<sup>5</sup> | 1.44×10<sup>4</sup> | - | 4.14×10<sup>3</sup> | 2.07×10<sup>6</sup> | 7.56×10<sup>3</sup> | - | - | 1.548×10<sup>5</sup> | - | 5.94×10<sup>3</sup> |
| Sample 5 | 1.584×10<sup>6</sup> | 4.14×10<sup>3</sup> | 8.28×10<sup>3</sup> | 3.708×10<sup>4</sup> | 3.024×10<sup>4</sup> | - | 9.54×10<sup>3</sup> | - | - | - | - | - | 2.6316×10<sup>6</sup> | 4.86×10<sup>3</sup> |
| Sample 6 | - | - | - | - | - | 4.554×10<sup>4</sup> | - | - | - | - | 3.492×10<sup>4</sup> | 2.16×10<sup>3</sup> | 1.044×10<sup>4</sup> | 4.68×10<sup>3</sup> |
| Sample 7 | - | - | - | - | 1.98×10<sup>3</sup> | 2.61×10<sup>4</sup> | 6.48×10<sup>3</sup> | - | - | - | - | 7.02×10<sup>3</sup> | - | 6.768×10<sup>4</sup> |
| Sample 8 | - | - | - | - | 1.566×10<sup>4</sup> | 1.926×10<sup>4</sup> | - | - | - | - | - | 1.98×10<sup>3</sup> | 1.296×10<sup>4</sup> | - |
| Sample 9 | - | - | - | - | - | 1.08×10<sup>4</sup> | - | - | - | - | - | - | 5.994×10<sup>5</sup> | - |
|  | GPR |  |  |  |  |  |  | *Staphylococcus* spp. |  |  |  |  |  |  |
| Sample 1 | 4.14×10<sup>5</sup> | 1.422×10<sup>4</sup> | 1.656×10<sup>4</sup> | 1.62×10<sup>4</sup> | 3.276×10<sup>4</sup> | 2.5488×10<sup>6</sup> | - | - | - | - | - | - | - | 3.1014×10<sup>5</sup> |
| Sample 2 | - | 1.098×10<sup>4</sup> | 5.004×10<sup>4</sup> | 1.53×10<sup>4</sup> | 6.498×10<sup>4</sup> | - | - | 1.62×10<sup>6</sup> | - | - | - | - | 1.8576×10<sup>6</sup> | - |
| Sample 3 | - | 8.82×10<sup>3</sup> | 2.88×10<sup>4</sup> | 9.18×10<sup>3</sup> | 9.216×10<sup>5</sup> | - | - | 5.886×10<sup>6</sup> | - | - | 1.656×10<sup>4</sup> | - | - | 8.136×10<sup>4</sup> |
| Sample 4 | - | 1.26×10<sup>4</sup> | 4.5×10<sup>4</sup> | - | 6.876×10<sup>5</sup> | 2.7432×10<sup>6</sup> | 8.982×10<sup>4</sup> | 1.278×10<sup>6</sup> | - | - | 2.9376×10<sup>6</sup> | - | - | - |
| Sample 5 | 6.12×10<sup>6</sup> | 1.62×10<sup>3</sup> | 5.76×10<sup>3</sup> | 1.818×10<sup>4</sup> | - | - | 1.494×10<sup>4</sup> | - | 7.74×10<sup>3</sup> | 1.152×10<sup>4</sup> | - | 9.09×10<sup>4</sup> | 2.7×10<sup>4</sup> | - |
| Sample 6 | - | - | 6.66×10<sup>3</sup> | 3.42×10<sup>3</sup> | 3.492×10<sup>4</sup> | 3.24×10<sup>3</sup> | 3.348×10<sup>4</sup> | - | - | 3.6×10<sup>3</sup> | 2.088×10<sup>4</sup> | - | - | - |
| Sample 7 | - | - | - | - | 1.602×10<sup>4</sup> | - | 4.68×10<sup>4</sup> | - | - | - | - | 4.14×10<sup>3</sup> | 4.86×10<sup>3</sup> | - |
| Sample 8 | - | - | - | - | - | 2.772×10<sup>4</sup> | - | - | - | - | - | 1.03968×10<sup>6</sup> | 5.04×10<sup>3</sup> | - |
| Sample 9 | - | - | - | - | - | 4.824×10<sup>5</sup> | - | - | - | - | - | - | 4.086×10<sup>5</sup> | - |
|  | *Chryseobacterium* spp. |  |  |  |  |  |  |  |  |  |  |  |  |  |
| Sample 1 | - | - | - | - | - | - | - |  |  |  |  |  |  |  |
| Sample 2 | - | 5.94×10<sup>3</sup> | 3.78×10<sup>3</sup> | - | - | - | - |  |  |  |  |  |  |  |
| Sample 3 | - | 7.2×10<sup>4</sup> | 1.8×10<sup>4</sup> | - | 3.834×10<sup>5</sup> | - | - |  |  |  |  |  |  |  |
| Sample 4 | - | 8.64×10<sup>3</sup> | 2.34×10<sup>4</sup> | - | - | - | 2.952×10<sup>4</sup> |  |  |  |  |  |  |  |
| Sample 5 | - | 1.188×10<sup>4</sup> | - | - | 3.456×10<sup>4</sup> | - | - |  |  |  |  |  |  |  |
| Sample 6 | - | - | - | - | 7.02×10<sup>3</sup> | - | 6.3×10<sup>3</sup> |  |  |  |  |  |  |  |
| Sample 7 | - | - | - | - | - | 1.296×10<sup>4</sup> | 4.14×10<sup>3</sup> |  |  |  |  |  |  |  |
| Sample 8 | - | - | - | - | - | - | - |  |  |  |  |  |  |  |
| Sample 9 | - | - | - | - | - | - | - |  |  |  |  |  |  |  |

GPR: Gram-positive rods, CFU: Colony Forming Units; -: absence of visible bacterial growth

### Bacteria susceptibility to antibiotics

Susceptibility tests to antibiotics were performed with 169 bacterial colony morphotypes (the distribution of numbers of bacterial colony morphotypes found is presented in the S1 table). These tests revealed several cases of antibiotic multidrug-resistance, with isolates from agricultural plots more frequently expressing resistant phenotypes than those from control plot (Table 3). The most effective antibiotics were Levofloxacin, Imipenem, Gentamicin and Ciprofloxacin.

**Table 3.** Antibiotic susceptibility profile.

| Antibiotics | Farm plots |  |  |  |  |  |  |  |  |  |  |  |  |  |  | Control plot |  |  |  |  |  |  |  |  |  |  |  |
| --- | --- | --- | --- | --- | --- | --- | --- | --- | --- | --- | --- | --- | --- | --- | --- | --- | --- | --- | --- | --- | --- | --- | --- | --- | --- | --- | --- |
|  | *Aeromonas* spp. |  |  | *Chryseobacterium* spp. |  |  | *Pseudomonas* spp. |  |  | *Staphylococcus* spp. |  |  | GPR |  |  | *Aeromonas* spp. |  |  | *Pseudomonas* spp. |  |  | *Staphylococcus* spp. |  |  | GPR |  |  |
|  | S | SHP | R | S | SFP | R | S | SHP | R | S | SHP | R | S | SHP | R | S | SHP | R | S | SFP | R | S | SHP | R | S | SHP | R |
| AMX | 26 | 0 | 74 | 14 | 0 | 86 | - | - | - | - | - | - | 56 | 6 | 38 | 17 | 0 | 83 | - | - | - | - | - | - | 0 | 0 | 100 |
| AMC | 41 | 0 | 59 | 36 | 0 | 64 | - | - | - | - | - | - | 35 | 0 | 65 | 83 | 0 | 17 | - | - | - | - | - | - | 67 | 0 | 33 |
| FOX | 46 | 18 | 36 | 57 | 7 | 36 | - | - | - | 17 | 0 | 83 | 73 | 13 | 13 | 50 | 0 | 50 | - | - | - | 40 | 0 | 60 | 67 | 0 | 33 |
| CRO | 15 | 3 | 82 | 21 | 0 | 79 | - | - | - | - | - | - | 23 | 0 | 77 | 83 | 0 | 17 | - | - | - | - | - | - | 67 | 0 | 33 |
| FEP | 5 | 5 | 90 | 21 | 0 | 79 | 21 | 0 | 79 | - | - | - | 19 | 0 | 81 | 17 | 17 | 67 | 50 | 25 | 25 | - | - | - | 100 | 0 | 0 |
| ATM | 21 | 5 | 74 | 29 | 0 | 71 | 25 | 0 | 75 | - | - | - | - | - | - | 83 | 0 | 17 | 50 | 0 | 50 | - | - | - | - | - | - |
| IMP | 62 | 3 | 36 | 64 | 7 | 29 | 50 | 4 | 46 | - | - | - | 29 | 2 | 69 | 83 | 0 | 17 | 75 | 0 | 25 | - | - | - | 100 | 0 | 0 |
| CIP | 62 | 15 | 23 | 71 | 0 | 29 | 50 | 14 | 36 | 6 | 61 | 33 | 69 | 10 | 21 | 100 | 0 | 0 | 100 | 0 | 0 | 20 | 80 | 0 | 33 | 67 | 0 |
| NOR | 54 | 0 | 46 | 57 | 0 | 43 | - | - | - | 50 | 0 | 50 | - | - | - | 100 | 0 | 0 | - | - | - | 100 | 0 | 0 | - | - | - |
| LEV | 54 | 18 | 28 | 64 | 7 | 29 | 68 | 7 | 25 | 78 | 11 | 11 | 92 | 4 | 4 | 100 | 0 | 0 | 100 | 0 | 0 | 80 | 20 | 0 | 67 | 33 | 0 |
| GEN | 46 | 0 | 54 | 43 | 0 | 57 | - | - | - | 78 | 0 | 22 | - | - | - | 50 | 0 | 50 | - | - | - | 100 | 0 | 0 | - | - | - |
| SXT | 0 | 10 | 90 | 0 | 14 | 86 | - | - | - | 6 | 6 | 89 | 15 | 0 | 85 | 67 | 0 | 33 | - | - | - | 40 | 20 | 40 | 33 | 0 | 67 |
| TET | - | - | - | - | - | - | - | - | - | 28 | 11 | 61 | 48 | 10 | 42 | - | - | - | - | - | - | 80 | 20 | 0 | 67 | 33 | 0 |
| PEN | - | - | - | - | - | - | - | - | - | 0 | 0 | 100 | - | - | - | - | - | - | - | - | - | 0 | 0 | 100 | - | - | - |

GPR: Gram-positive rods, S: frequencies of susceptible isolates, SHP: frequencies of isolates susceptible at high posology, R: frequencies of resistant isolates, AMX: Amoxicillin (20 or 25 µg), AMC: Amoxicillin/Clavulanic acid (20/10 µg), ATM: Aztreonam (30 µg), CIP: Ciprofloxacin (5 µg), CRO: Ceftriaxone (30 µg), FEP: Cefepime (30 µg), FOX: Cefoxitin (30 µg), GEN: Gentamicin (10 µg), IMP: Imipenem (10 µg), LEV: Levofloxacin (5 µg), NOR: Norfloxacin (10 µg), PEN: Penicillin G (10 U), TET: Tetracycline (30 µg), SXT: Trimethoprim/Sulphamethoxazol (1.75/23.25 µg); - :not tested,

## Discussion

Data analysis from the present survey conducted in Mangoum’s farmlands, a neighborhood of the Foumbot municipality (Noun division, West Cameroon), primarily revealed that all farmers used animal manure and pesticides in crop production. Pesticides were applied every 3, 4 or 5 days, depending on the farmer will. Manures basically originated from avian and swine farms and applied before soil plowing and seeding. The most recent application dates prior to the present study were found between 2 and 6 weeks. This overall tendency to use fertilizers and pesticides resides in the logic that targets sustainable higher level production of good-quality crops in line with the ever-growing populations demands for better welfare that should couple with economic benefits for farmers. This could explain the relentless efforts in plant protection against pests and invaders in addition to plant growth supplements that is common in the study area. However, these practices corelate with higher risk of generating factors which are likely to promote selection of antibacterial resistance (ABR) and dissemination of antimicrobial resistance genes (ARG) [6-10]. The phenomena of ARG selection and spread, more obvious with bacteria may concern other prokaryotes within and amongst ecological niches, in connection with the microbial genome flexibility. Acknowledging that the genetic code is not only universal but degenerated as well, anticipating adverse effects on exposed eucaryotes is reasonable.

Major bacteria recovered were Gram-positive rods, *Aeromonas* spp., *Chryseobacterium* spp., *Pseudomonas* spp., and *Staphylococcus* spp., in subjected soil specimens. Known as endogenous environmental bacteria flora, their loads in the farmland soils were significantly lower than in the control plot. Since the visible difference between the control and farm soils is alleged through the absence or the presence of transformations and treatments made on these soils, data analysis further highlights that microbial populations are adversely affected by human activities, consistent with previous observations on agropastoral activities and microbial populations [14]. Otherwise, although useful for crop protection, pesticides do affect soil microflora beyond expectation ranges. Their repeated use maintains pressure that might cause evolution in the niche microbiota, altering thereby the inherent microbial soil characteristics in types and diversity [15,16]. These events are likely to effect in the long run the target soil’s mineralogy and the overall ‘‘soil health’’ subsequent to population evolution. If human activities can explain the difference in bacteria loads between the farms and control plot, inherent specificities of practices on each agricultural plot could explain certain differences in bacterial loads between the target agricultural plots. These specificities could have created variations in the effect of treatments and activities on microbial populations, consistent with above development on risk of microbiota alteration that may evolve irreversibly with sustainable pressure, an affect any future agricultural project. Other factors like the chemical and organic composition of soils, soil characteristics, abiotic and biotic factors might also explain these plot-specific variations in the recorded bacterial loads [17-19] in line with varied practices. Otherwise, harmonized agricultural policies guided by local-accompanying trained leaders are necessary. The present investigation basically revealed that application of manure and/or pesticides depended on the farmer will, without any reference though they claimed to be trained in agriculture. For instance, gloves and boots were rarely used (if ever), in addition to lack of clean water for onsite-bath.

The paucity of biosafety and biosecurity and an absence of knowledge regarding antimicrobial resistance further emphasize the need for capacity building. This should stand as a pre-requisite for a safer application of pesticides and animal manure, consistent with their likely adverse impact on exposed human and animal populations beyond alteration of the microbial flora.

With a glance on bacterial susceptibility, a reduce number on antibacterial agents used was effective on bacteria isolated from agricultural soils compared to those from the control plot. Globally, fluoroquinolones, Imipenem and Gentamicin were most effective on bacterial isolates. Like bacterial loads, this difference further highlights the impact that soil transformations and treatments can have on endogenous microbial populations.

Two factors can be pointed out as very likely to have contributed to the higher resistance rates in farmlands. The first one is the use of manure. On the farm soils investigated, the manure applied consisted of feces of farm animals that are known reservoirs of antibiotic residues, antibiotic-resistant germs and mobile genetic elements [24]. Accordingly, in the presence of manure, soil bacteria are exposed to selective pressures caused by antibiotic-resistant strains and genes that have been selected in animal digestive tracts and/or in the farm environment on one hand, and to the pressure caused by antibiotic residues contained in it on the other. These multivariate stresses combine and exacerbate acquisition rates of resistance genes and thus, the emergence of more resistant bacteria populations [20-23]. This development agrees with findings from previous survey conducted in animal farms in Cameroon (West [24,25], Littoral [26], South [27]), which reported high levels of antibiotic-resistant bacteria in animal feces and in some items within the farm premises like animal feed and drinking water. Also, the clinical category profiles of isolates from these previous studies overlap with those recorded during the present one. This overlapping tendency might highlight the links between animal farms and plant farms. This view should be taken into account in all initiatives that focus accurately antimicrobial resistance according to the One Health principles (multisite, multidisciplinary, gender-related, holistic view, to name a few) in Cameroon.

Pesticides application is the second engine that likely contributed to antibiotic resistance selection. In fact, bacteria can express resistance to antibiotics when they are exposed to other biocides by cross-selection, co-selection, and phenotype change. Also, indirect promotion of less susceptible microbial sub-population upon exposure to biocides occurs through several mechanisms including activation of *SOS* response, repair of DNA alteration induced which ultimately result in microbial fitness [28] and altered environmental microbial diversity.

Additionally, the results recorded indicate that these farmers are exposed to a diversity of multidrug-resistant bacteria, likely etiologies of infections. Consequently, they could be acting as disseminators of these bacteria, driving therefore the spread of ARB and ARG into their communities. The paucity in biosecurity-biosafety further exacerbates this role of transporter. Based on current data and finings, however, it is not possible to anticipate which driver (manure or pesticides) was most effective in selecting resistance traits, and to what extent.

Higher rates of ARB in farm soils justify the need for amplified investigations on ARG in crop production environments in order to track and control ARB and ARG dissemination in vulnerable settings and communities in Cameroon. The present survey was conducted during the raining season. The rainfall was identified by previous authors as a mobile reservoir facilitating the transmission and proliferation of ARG, and enhancing bacterial resistance and the abundance of ARG in soil [29]. Then, future investigations should include meteorological parameters such as seasonal changes in the study design to better track the selection and diffusion of ARB and ARG in Cameroon’s agricultural sector.

This links that bring together West Cameroon’s agricultural farms, animal farms and exposed communities further reinforces the need to emphasize the One Health paradigm from the holistic point of view. In this, current data provide support to advocate One Health activities such as raising farmers’ awareness and improving policies that focus issues associated with antimicrobial resistance for global welfare.

## Conclusion

The present investigation revealed that the “agricultural plot” characteristic had a great effect on bacteria from soil microbial flora of agricultural farms in Mangoum locality, West Cameroon. The application of pesticides and manure in plant farm was very likely associates with selection of antibiotic-resistant bacterial strains. Rates of antibiotic-resistant bacteria were higher in farm soils. Also, based on current data analysis, it wasn’t possible to specify which substrate (manure or pesticides) was most effective in selecting the resistance traits observed. Overall, the most effective antibiotics were Levofloxacin, Imipenem, Gentamicin and Ciprofloxacin. Agreeing with previous surveys conducted on animal farms with similar environmental conditions, these findings could provide support to sustainable orientation of policies regarding control of antimicrobial resistance in Cameroon from the One Health perspective.

## Acknowledgements

The authors thank the “Université des Montagnes” and its teaching hospital for the support during this survey. The authors thank the farm owners for the availability and cooperation.

## Supporting information

**S1 table. Number of bacterial colony morphotypes found**

## Notes

### Competing Interest Statement

The authors have declared no competing interest.

